# Repetitive Spreading Depolarization induces gene expression changes related to synaptic plasticity and neuroprotective pathways

**DOI:** 10.1101/2023.02.27.530317

**Authors:** Michela Dell’Orco, Jordan E. Weisend, Nora I. Perrone-Bizzozero, Andrew P. Carlson, Russell A. Morton, David N Linsenbardt, C. William Shuttleworth

## Abstract

Spreading depolarization (SD) is a slowly propagating wave of profound depolarization that sweeps through cortical tissue. While much emphasis has been placed on the damaging consequences of SD, there is uncertainty surrounding the potential activation of beneficial pathways such as cell survival and plasticity. The present study used unbiased assessments of gene expression to evaluate that compensatory and repair mechanisms could be recruited following SD, regardless of the induction method, which prior to this work had not been assessed. We also tested assumptions of appropriate controls and the spatial extent of expression changes that are important for *in vivo* SD models. SD clusters were induced with either KCl focal application or optogenetic stimulation in healthy mice. Cortical RNA was extracted and sequenced to identify differentially expressed genes (DEGs). SDs using both induction methods significantly upregulated 16 genes (versus sham animals) that included the cell proliferation-related genes FOS, JUN, and DUSP6, the plasticity-related genes ARC and HOMER1, and the inflammation-related genes PTGS2, EGR2, and NR4A1. The contralateral hemisphere is commonly used as control tissue for DEG studies, but its activity could be modified by near-global disruption of activity in the adjacent brain. We found 21 upregulated genes when comparing SD-involved cortex versus tissue from the contralateral hemisphere of the same animals. Interestingly, there was almost complete overlap (21/16) with the DEGs identified using sham controls. Neuronal activity also differs in SD initiation zones, where sustained global depolarization is required to initiate propagating events. We found that gene expression varied as a function of the distance from the SD initiation site, with greater expression differences observed in regions further away. Functional and pathway enrichment analyses identified axonogenesis, branching, neuritogenesis, and dendritic growth as significantly enriched in overlapping DEGs. Increased expression of SD-induced genes was also associated with predicted inhibition of pathways associated with cell death, and apoptosis. These results identify novel biological pathways that could be involved in plasticity and/or circuit modification in brain tissue impacted by SD. These results also identify novel functional targets that could be tested to determine potential roles in recovery and survival of peri-infarct tissues.

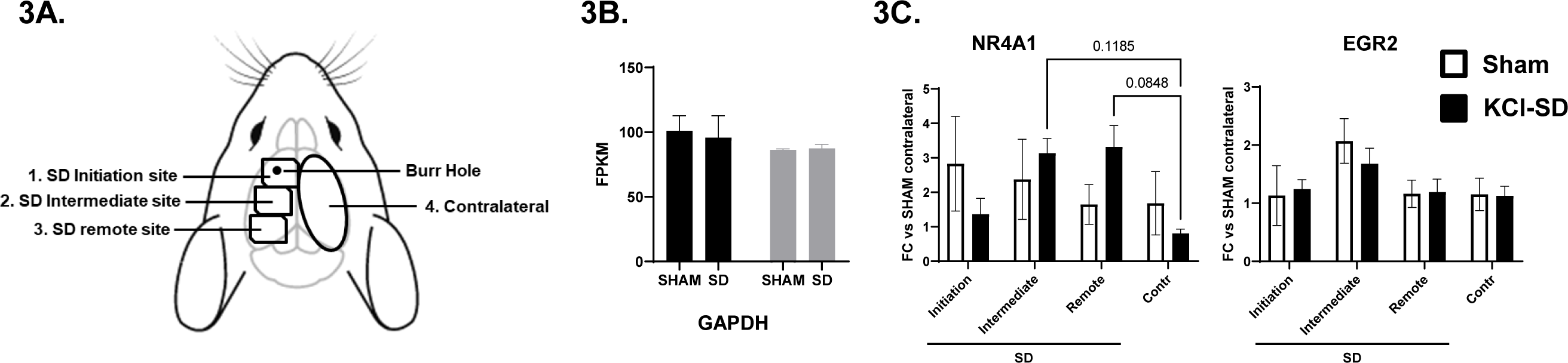

## 1 Introduction

Spreading Depolarization (SD) is a slowly propagating wave of extended depolarization that is followed by transient suppression of neural activity, first described by Leao (Leao, 1944). SD is implicated as a mechanism of migraine with aura (Charles and Brennan, 2009; Pietrobon and Moskowitz, 2014) and otherwise healthy brain is able to fully recover from SD (Nedergaard and Hansen, 1988). However, the long duration of depolarization (up to a minute or more) can lead to severe energy depletion (Lindquist and Shuttleworth, 2014; Dreier et al., 2017), particularly when coupled with neurovascular disruptions in ischemic brain (Dreier, 2011; Ayata and Lauritzen, 2015; Hartings et al., 2017b). In addition, transient extracellular glutamate surges caused by SD can be sufficient to cause neuronal injury in metabolically compromised brain tissue (Aiba et al., 2012; Hinzman et al., 2015). Recent work has identified SDs as contributors to tissue loss following stroke, trauma, and other acute brain injuries (Lauritzen et al., 2011; Dreier et al., 2017) and initial efforts have begun targeting SD in efforts to improve outcomes in brain injury (Carlson et al., 2018; Helbok et al., 2020).

In contrast to the large body of literature characterizing electrophysiological and vascular responses to SD, only a relatively small number of studies have examined SD-induced gene expression changes and their relationship to outcomes in different experimental and clinical conditions (Urbach et al., 2006; Sintas et al., 2017). Initial studies identified increases in brain derived neurotrophic factor (BDNF), c-Fos and heat shock protein 70 (HSP70) (Kokaia et al., 1993; Matsushima et al., 1996; Karikó et al., 1998; Dietrich et al., 2000; Rangel et al., 2001). Subsequent studies showed increased expression of pro-inflammatory and apoptotic genes, including tumor necrosis factor (TNF), C-C motif chemokine ligand 2 (CCL2), interleukin-1 (IL-1) and -6, and cyclooxygenase-2 (COX-2), and B-cell lymphoma 2 (BCL-2) (Jander et al., 2001; Kaido et al., 2012; Eising et al., 2017; Takizawa et al., 2020). While most of these studies focused on a predetermined set of genes, a microarray study applied gene ontology (GO) and pathway analysis (PA) to examine the effect of SDs in rats (Sintas et al., 2017). This study identified upregulation of genes in pathways linked to oxidative stress and cellular injury, but less is known of other cellular adaptations that may occur. A lack of comprehensive transcriptomic analysis has limited the understanding of potentially beneficial pathways that could be activated following SD.

In the current study, we provide the first unbiased RNAseq analysis following SD. To increase generalizability, we combined data from the two most common methods of SD initiation (focal depolarization with either topical KCl administration or optogenetic stimulation) and generated clusters of SDs that are analogous to patterns of SDs that have been recorded from injured human brain. We also examined two key issues specific this type of study in SD models; 1) appropriate control tissues and 2) relative contribution of initiation and propagation sites to gene expression changes. Finally, we applied pathway analysis approaches to uncover candidate networks and novel genes altered by SD. The results identify the upregulation of multiple molecular pathways in the propagation zones of SD that could regulate potentially beneficial effects, such as neuritogenesis and synaptic plasticity.

## 2 Materials and Methods

### 2.1 Animal model

All experimental procedures were performed in accordance with the National Institutes of Health Guide for Care and Use of Laboratory Animals and were approved by the University of New Mexico Health Sciences Center Institutional Animal Care and Use Committee (IACUC). SDs were induced repetitively (4 SDs in 2 hours at 30 min intervals) in healthy 7-8-week-old female mice and SDs were confirmed with intrinsic optical signal imaging (Video 1, 2, and Supplementary Figure 1). SDs were induced in 6 mice with either focal application of KCl through a burr hole in C57Bl6J mice (n=3) or with optogenetic stimulation (3 mW for 20 seconds) in Thy1-ChR2-YFP positive mice (n=3). Thy1-ChR2-YFP breeders were initially purchased from The Jackson Laboratory (B6.Cg-Tg(Thy1-COP4/EYFP)18Gfng/J, Strain #:007612, RRID:IMSR_JAX:007612) and crossed to maintain the line, as previously described (Arenkiel et al., 2007). With either stimulus, evoked SDs propagated throughout the neocortex ipsilateral to the stimulus, without crossing to the contralateral hemisphere **(Supplementary Figure 1, Video 1 and 2).** A matched number of sham controls was used with focal NaCl administration through a burrhole, in C57Bl/6 mice (n=3), or 0.4 mW for 20 seconds in Thy1-ChR2-YFP positive mice (n=3). Thirty minutes after the last SD, the anesthetized animal was decapitated and the brain extracted, washed with ice cold PBS and kept on ice.

### 2.2 RNA extraction and qRT-PCR validation

The contralateral and ipsilateral cortices were rapidly dissected from underlying striatum and the hippocampus and total RNA was extracted using TRIzol (Invitrogen, ThermoFisher, Waltham, MA. USA). Extractions were either from intact ipsilateral or contralateral cortices (Figures 2-7) or from regions ∼18 mm^2^ rapidly micro-dissected on ice (Figure 6). Where indicated, microdissections were made from 3 different regions of the ipsilateral hemisphere: 1) SD initiation site; 2) intermediate site > 3mm from initiation, 3) remote site >5 mm from initiation (Supplementary Figure 3). All collected tissues were then flash frozen and kept at –80°C until RNA extraction. RNA was quantified using Qubit (Bio-Rad) and its quality was determined using a NanoDrop 1000 (ThermoFisher), using absorbance at 260, 280 and 230 nm. Aliquots of 2 µg RNA were sent to Arraystar, Inc. for Illumina paired-end RNAseq as described below. To validate gene expression from our RNAseq data set, separate aliquots containing 1 μg of total RNA were reverse transcribed using SuperScript II reverse transcriptase (Life Technologies) following the manufacturer’s protocol and qPCR carried out with a CFX96 Touch Real-Time PCR Detection System using SYBR Green mix (Life Technologies). Primer sequences for qRT-PCR for mouse mRNAs were obtained from PrimerBank (https://pga.mgh.harvard.edu/primerbank/). The GAPDH gene (FW TGTGATGGGTGTGAACCACGAGAA, RV GAGCCCTTCCACAATGCCAAAGTT) was used to normalize values. As shown in Supplementary Figure 3, GAPDH expression itself was unchanged by SD with either KCl or optogenetic induction of SD. The relative expression of target genes was determined using the comparative 2−ΔCt method (Livak and Schmittgen, 2001).

Values were expressed as means ± SEM. Statistical analysis was performed using GraphPad Prism version 9 (La Jolla, CA, USA). The data were analyzed by unpaired t-test followed by Welch’s correction or analysis of variance (ANOVA) followed by Dunnett’s Multiple Comparison tests. Differences were considered statistically significant when p values were < 0.05 (*p<0.05, **p<0.01, ***p<0.001).

### 2.3 RNA sequencing

RNAseq analysis was performed by Arraystar Inc. (Rockville, MD). Total RNA (2 µg) was used to prepare the sequencing library in the following steps: total RNA was enriched by oligo (dT) magnetic beads (rRNA removed); RNAseq library was prepared using KAPA Stranded RNASeq Library Prep Kit (Illumina), which incorporates dUTP into the second cDNA strand and renders the RNAseq library strand-specific. The completed libraries were qualified with an Agilent 2100 Bioanalyzer and quantified by using the absolute quantification qPCR method. To sequence the libraries on an Illumina NovaSeq 6000 instrument, the barcoded libraries were mixed, denatured to single stranded DNA in NaOH, captured on Illumina flow cell, amplified *in situ,* and subsequently sequenced for 150 cycles for both ends on an Illumina NovaSeq 6000 instrument. Sequence quality was examined using the FastQC software. The trimmed reads (trimmed 5’, 3’-adaptor bases using cutadapt) were aligned to a reference genome (Mouse genome GRCm38) using Hisat2 software.

### 2.4 Differential expression analysis

The expression levels of known genes and transcripts were calculated using the StringTie and Ballgown software package (Pertea et al., 2016) and expressed as fragments per kilobase of transcript per million mapped reads (FPKM). The number of identified genes and transcripts per group was calculated based on the mean of FPKM in group > 0.5. Differentially expressed gene (DEG) and transcript analyses were performed using an unpaired t-test followed by Welch’s correction; a fold change (FC) threshold of 1.25 and p value < 0.05 were used together to classify genes and transcripts as differentially expressed. We subsequently performed DESeq2 analysis (Ge et al., 2018) with a false discovery rate (FDR) correction cutoff of 0.01 and FC >1.5.

### 2.5 Ingenuity Pathway Analysis

Predicted molecular functions and biological pathways enriched in genes differentially expressed after SD were identified using Ingenuity Pathway software (IPA, Content version: 62089861 (Release Date: 2021-02-17), Qiagen, https://www.qiagenbioinformatics.com/products/ingenuity-pathway-analysis, (Krämer et al., 2014)). The analysis was set to filter nervous system expressed genes only; reference set: Ingenuity Knowledge Base (Genes Only); and including direct and indirect relationships.

## 3 Results

### 3.1 Identification of gene expression induced by repetitive spreading depolarizations

Healthy female mice were subjected to multiple SDs (4 over a period of 2 hours) which were generated to match clusters of SDs in clinical settings (Dreier et al., 2017) This procedure has previously been shown to induce robust transcriptional changes (Urbach et al., 2006; Sintas et al., 2017; Takizawa et al., 2020). Thirty minutes after the last SD or sham stimulus, the neocortex was removed, and RNA extracted from tissue subjected to SD or shams for subsequent RNA sequencing (see Materials and Methods and **Supplementary Figure 1**).

Both focal KCl and optogenetic stimulation are utilized in studies of SD in brain injury, and thus to increase generalizability of this initial study of SD regulation of target genes, data from these two SD-stimulation methods were combined for differential expression (DE) analysis. For the combined DE analysis, we performed unpaired t-tests followed by Welch’s correction on the mapped 12944 genes. We identified 102 significantly differentially expressed genes (DEGs) **(Figure 1A, black, FC > 1.25, p value <0.05, n=6, Supplementary Table 1).** We subsequently performed DESeq2 analysis (Ge et al., 2018) and identified 16 genes that were also significant after false discovery rate correction (FDR <0.01, FC>1.5) when comparing the SD group with the sham tissues (**Figure 1A, red, FDR <0.01, FC** >1.5, n=6, Supplementary Table 1).

**Figure 1.**
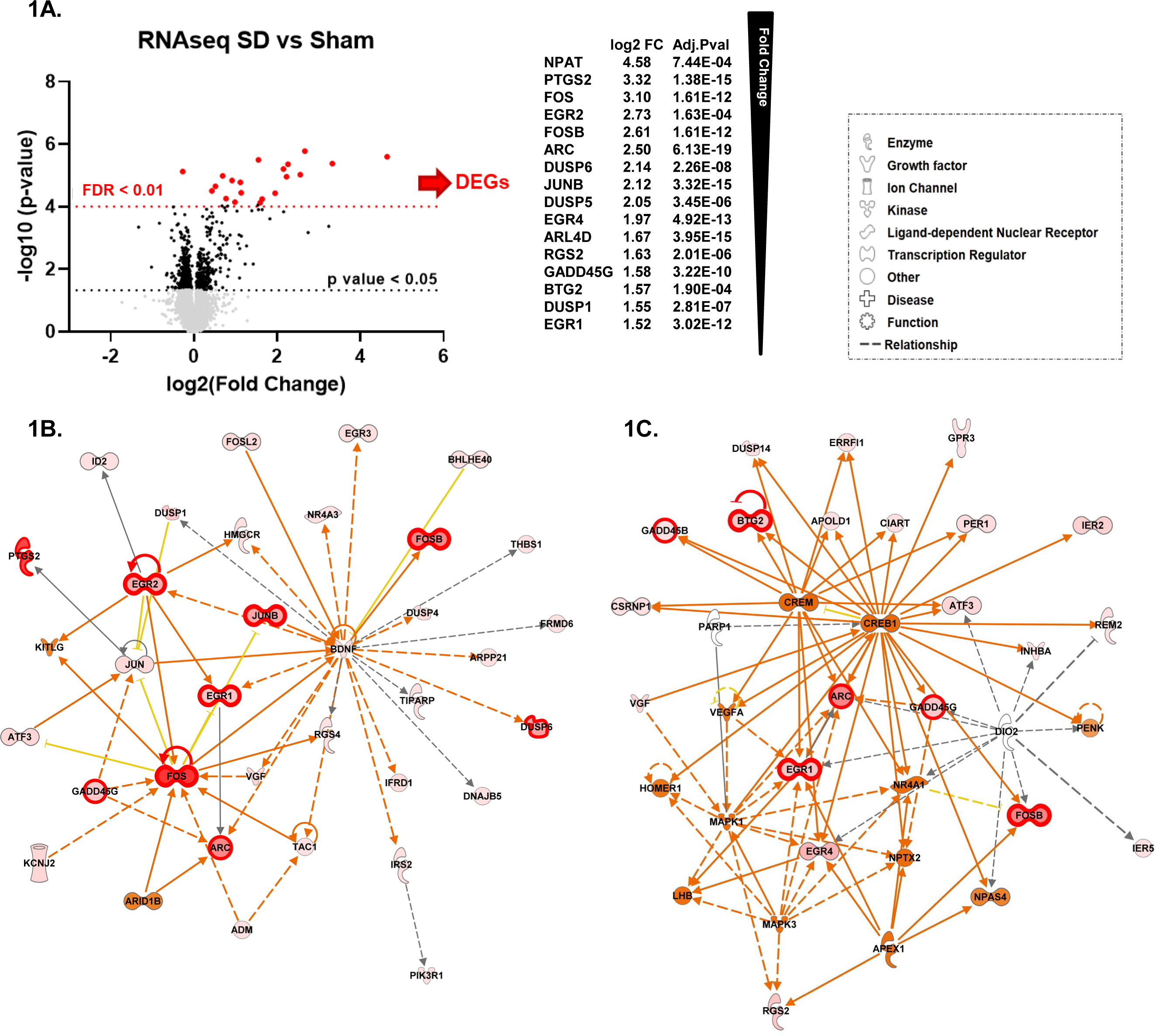
Repetitive SD increases levels of genes related to nervous system development and function. **A)** Differential Expression (DE) analysis. We performed unpaired t-test followed by Welch’s correction; significant genes (FC > 1.25, p value <0.05, n=6) are represented in black. Consequently, we performed DESeq2 analysis (Ge, Son, and Yao 2018); genes that were also significant after false discovery rate correction (FDR <0.01, n=6) are represented in red. **B) and C)** top molecular networks (score 63 and 32) generated by IPA analysis of DEGs upregulated in SD animals vs sham. The intensity of the color red for each molecule is proportional to the FC of the gene. Genes with FDR<0.01 are highlighted with a thicker red outline. Full arrows indicate a direct relationship between molecules, while dotted arrows indicate an indirect relationship between genes. Orange arrows and molecules indicate the predicted activation of a gene, while yellow indicates predicted inhibition. **D)** Top diseases and functions determined from the IPA analysis.

Consistent with previous studies (Kokaia et al., 1993; Dietrich et al., 2000; Rangel et al., 2001; Kaido et al., 2012), we found significant changes in immediate early genes ARC, FOS and JUN proto-oncogenes, and early growth response (EGR) genes. Other DEGs with significantly increased levels included adrenoceptor beta 1 (ADRB1), dual specificity phosphatase (DUSP) gene family, the nuclear receptor subfamily 4 group A member 1 (NR4A1), and prostaglandin-endoperoxide synthase 2 (PTGS2)

### 3.2 Molecular networks regulated by repetitive SDs

We next used Ingenuity Pathway Analysis (IPA) to identify molecular networks and biological pathways regulated by DEGs. This information was then used to obtain a prediction of physiological and pathological brain processes altered by SD. Enriched pathway analysis on FDR significant genes showed that genes upregulated by SD are involved in nervous system development and function, tissue morphology and development. We also found significant enrichment in genes involved in cell death and survival and cell morphology (**Supplemenraty Table 2**)

To achieve a more extensive enrichment analysis we then expanded IPA pathway analysis to genes with p value< 0.05 and FC> 1.25. **Figure 1B** and **C** show the top two scored networks (score 63 and 35) generated by IPA with the combined DEGs upregulated in the SD group as compared to sham animals. The intensity of the color red for each molecule is proportional to the fold change (FC) of the labeled gene. Genes with FDR< 0.01 are also highlighted with thicker red outlines. Full arrows indicate a direct relationship between molecules, while dotted arrows indicate a predicted indirect relationship. Orange arrows and molecules indicate the predicted activation of a gene, while yellow indicates predicted inhibition. These networks implicated SD target genes associated with cell proliferation and synaptic activity, including cAMP responsive element binding protein 1 (CREB1), mitogen-activated protein kinase 1 and 3 (MAPK1 and MAPK3), and HOMER1 (**Figure 1C**).

Predictions of SD-induced changes associated with known disease states and functions are shown in **Table 1** and they include: neurological diseases, psychological disorders, nervous system development and function, tissue morphology, and tissue development. The complete list of diseases and functions and the genes associated with each pathway are reported in **Supplementary Table 2**. **Figure 1D** also shows the number of genes associated with each pathway and associated p value range. They are shown together so that different strengths of the network elements can be compared. A complete list of networks and associated DEGs is reported in **Supplementary Table 3.**

**Table 1.** Predictions of SD-induced changes in associated nervous system diseases and functions, in SD vs Sham Animals.

| Disease and Disorders | p-value range | # Molecules |
| --- | --- | --- |
| Neurological Diseases | 3.69E-02 - 7.77E-09 | 42 |
| Organismal Injury and Abnormalities | 3.69E-02 - 6.10E-09 | 43 |
| Psychological Disorders | 2.65E-02 - 6.10E-09 | 29 |
| Hereditary Disorders | 2.65E-02 - 6.10E-09 | 18 |
| Skeletal and Muscular Disorders | 2.65E-02 - 2.70E-08 | 17 |
| Physiological System Development and Functions | p-value range | # Molecules |
| Nervous System Development and Function | 3.69E-02 - 7.62E-07 | 51 |
| Tissue Morphology | 3.69E-02 - 4.54E-08 | 28 |
| Tissue Development | 3.46E-02 - 7.62E-07 | 40 |
| Organismal Development | 3.46E-02 - 4.43E-06 | 37 |
| Behavior | 3.17E-02 - 1.11E-05 | 12 |

Figure 2 illustrates pathways related to cellular functions induced by repetitive spreading depolarization. The top common pathways identified include: cellular assembly and organization, cellular function and maintenance, and cellular development and proliferation (**Supplementary Table 4**). The heatmaps in Figure 2A and B show the p value and the activation z score for each pathway, and a graphical representation of the cellular functions and genes involved, respectively. As shown in Figure 2, each gene expression FC is proportional to their color intensity; orange arrows and pathways indicate a predicted activation, whereas blue indicates increased levels of genes which lead to inhibition of the pathway; genes with FDR< 0.01 are also highlighted with thicker red outlines. IPA analysis predicted an activation of axonogenesis, axon branching, dendritic growth and branching and growth of neurites, based on increased levels of genes such as VGF, FOS, PTGS2, JUN, BDNF, gamma-aminobutyric acid type A receptor subunit beta3 (GABRB3), activating transcription factor 3 (ATF3), and cyclin dependent kinase like 3 (CDKL3). We also found predicted activation of pathways regulating neuronal development, long-term potentiation, and neurotransmission due to increased expression of genes such as FOS, PTGS2, EGR2, BDNF, JUN, KCNJ2, ARC, and HOMER1.

**Figure 2.**
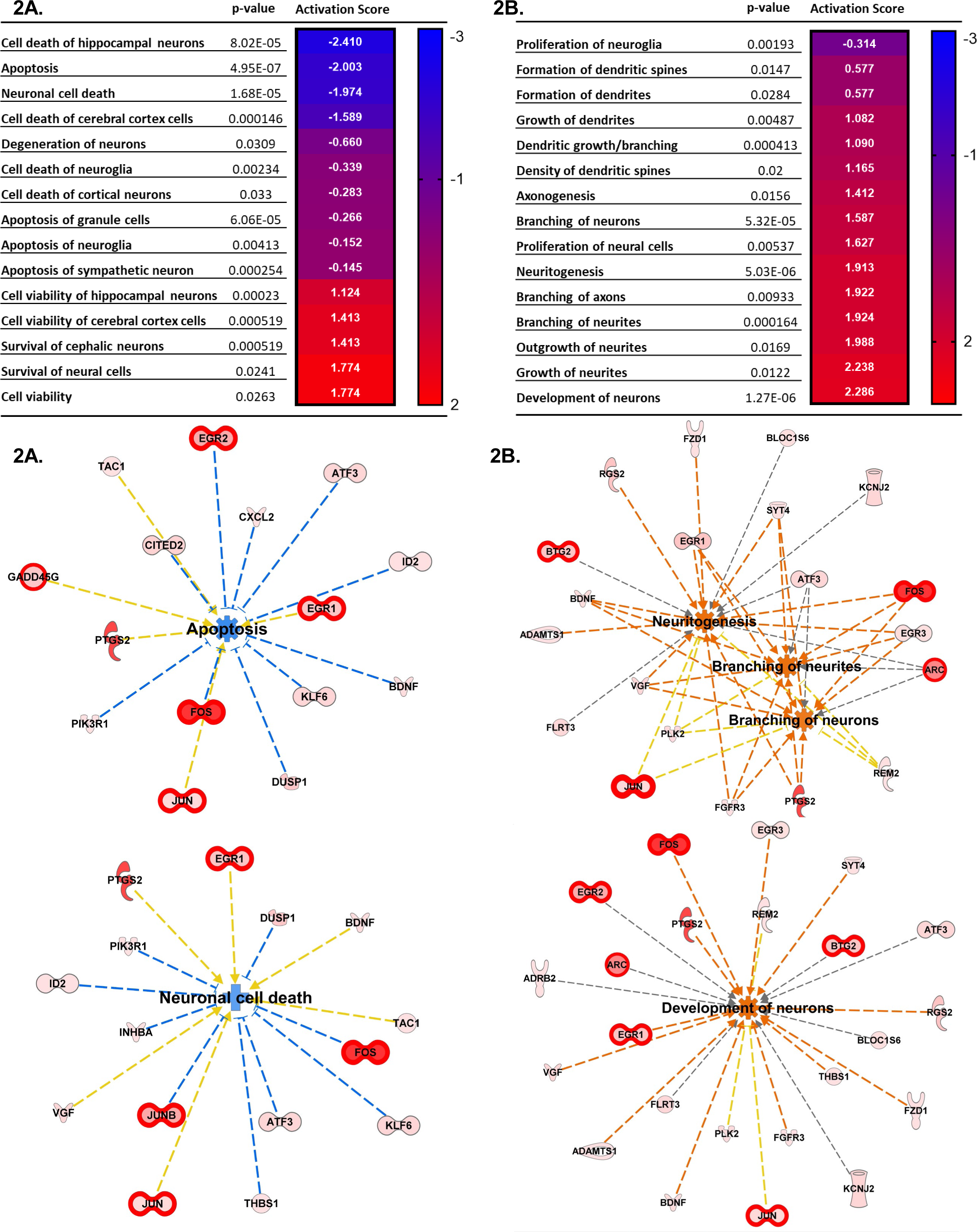
Top cellular pathways activated by SD. **A)** Neural death and survival pathways. **B)** Cellular proliferation and growth. Top cellular functions generated by IPA analysis of DEGs upregulated in SD animals vs sham. The heatmaps show the p value and the activation score for each pathway. **A’) and B’)** Graphical representations of the networks and the genes involved in A and B. The intensity of the color red for each molecule is proportional to the FC of the gene. Orange arrows indicate a positive control of the gene on the related pathway, and the pathway’s name highlighted in orange indicates its activation. Genes with FDR< 0.01 are also highlighted with thicker red outlines. Apoptosis z score -1.197, neuronal cell death z score -0.903, neuritogenesis z score +2.048, branching of neurites z score +1.924, branching of axons z score +1.922, development of neurons z score +2.224.

Interestingly, we also found predicted inhibition of neuronal cell death (z score -0.903), apoptosis (z score -1.197) (Figure 2A), together with increased development of neurons (z score +2.224) survival of cerebral cortex cells (z score +1.774), and cell viability (z score +1.413) (Figure 2B), due to increased expression of genes such as FOS, BDNF, ATF3, JUNB, DUSP1, neuronal PAS domain protein 4 (NPAS4), nuclear factor interleukin 3 regulated (NFIL3), and SERPINE1.

### 3.3 Comparison between contralateral and ipsilateral hemispheres

Many studies utilize the contralateral neocortex as a convenient comparison tissue for DEG studies (Shen and Gundlach, 1998; Volobueva et al., 2023), as SD does not directly propagate across the corpus collosum. However, even if SD does not directly propagate through the contralateral cortex, the profound and widespread disruption of neuronal network activity produced by SD might be expected to have some impact on neuronal activity and gene expression in the contralateral hemisphere. The assumption that contralateral tissue is equivalent to tissue from sham animals for DE analysis has not previously been tested.

We first performed DE analysis between these 2 tissues (i.e., contralateral hemispheres in animals exposed to SD in the ipsilateral hemisphere vs. sham tissues). As shown in **Supplementary Figure 2** only 9 genes were significantly differently expressed after unpaired t-test followed by Welch’s correction, but none of were still significant after FDR correction, confirming a high degree of similarity between gene expression in these tissues.

We next performed the same analyses described above (Figure 1), but comparing tissue exposed to repetitive SDs with the contralateral cortex in the same animals, rather than sham animals. Of the 12893 mapped genes 149 were significant after unpaired t-test followed by Welch’s correction (Figure 3A, black, FC >1.25, p value <0.05, n=6, **Supplementary Table 1**), of these 21 were also significant after FDR correction (FDR <0.01, FC>1.5, n=6) (Figure 3A, red). IPA analysis confirmed a high similarity between the sham and contralateral cortex when used as control compared to the SD group. Figure 3B and 3C show the top scored networks generated from the SD vs contralateral DEGs list, genes with FDR< 0.01 are also highlighted with thicker red outlines. The top diseases and function are reported in **Table 2** and Supplementary Table 2 (compare with Figure 1). Likewise, Figure 4 shows cellular and molecular pathways that are comparable with DEG analysis with sham tissues (compared with Figure 2). These results imply that, at least at the level of gene expression changes after 2 hours, the contralateral hemisphere that is not directly invaded by SD can likely serve as an unaffected control.

**Figure 3.**
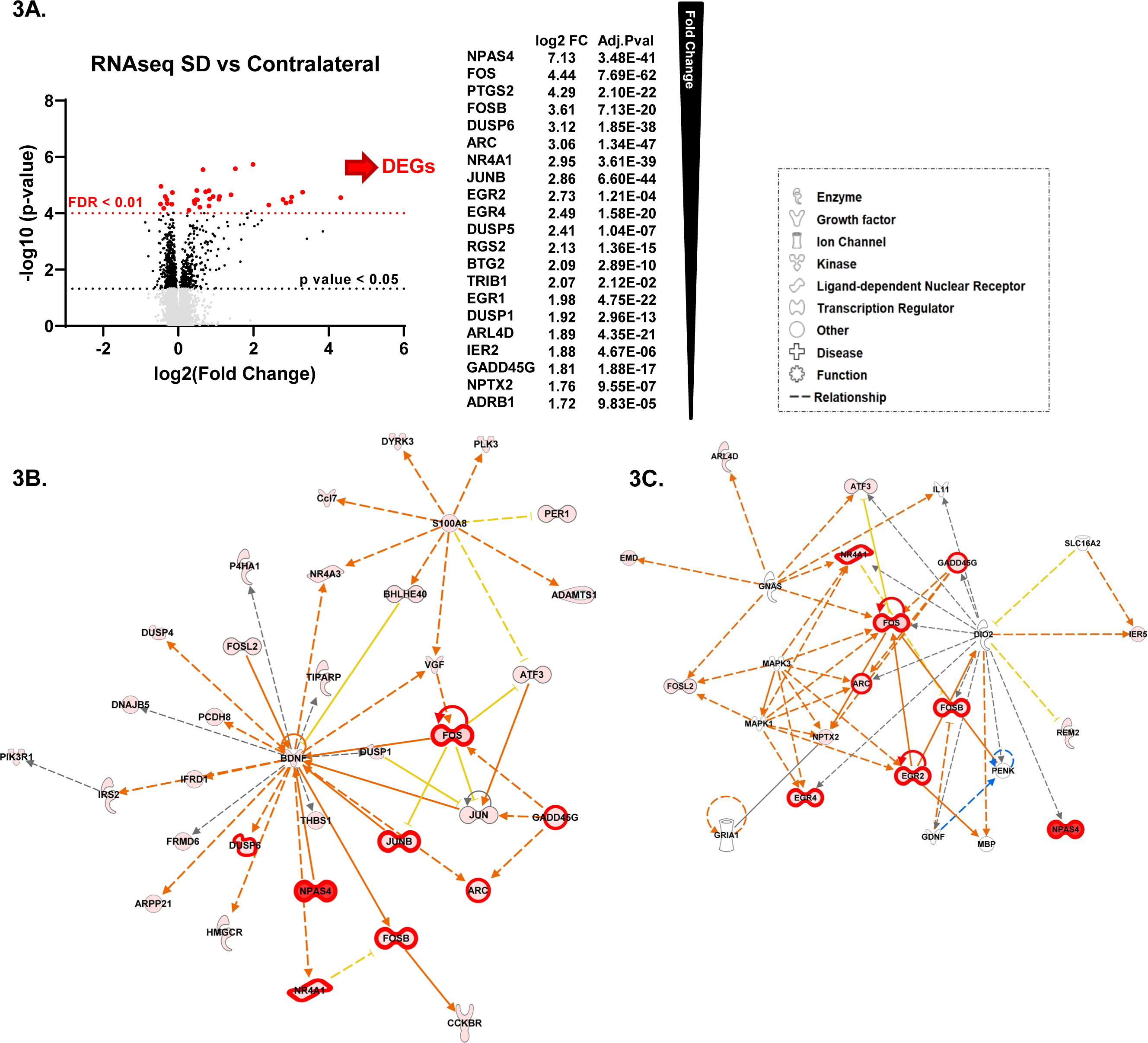
Sham animals and contralateral cortex are equivalent controls for unilateral SDs. **A**) Differential Expression (DE) analysis. We performed unpaired t-test followed by Welch’s correction; significant genes (FC > 1.25, p value <0.05, n=6) are represented in black. Subsequently, we performed DESeq2 analysis (Ge, Son, and Yao 2018); genes that were also significant after false discovery rate correction (FDR <0.01, n=6) are represented in red. **B**) top molecular networks (score 65 and 34) generated by IPA analysis of DEGs upregulated in SD animals vs contralateral. The intensity of the color red for each molecule is proportional to the FC of the gene. Full arrows indicate a direct relationship between molecules, while dotted arrows indicate an indirect relationship between genes. Orange arrows indicate the predicted activation of a gene, while yellow indicates predicted inhibition. Genes with FDR< 0.01 are also highlighted with thicker red outlines. **C**) Top diseases and functions determined from IPA analysis.

**Figure 4.**
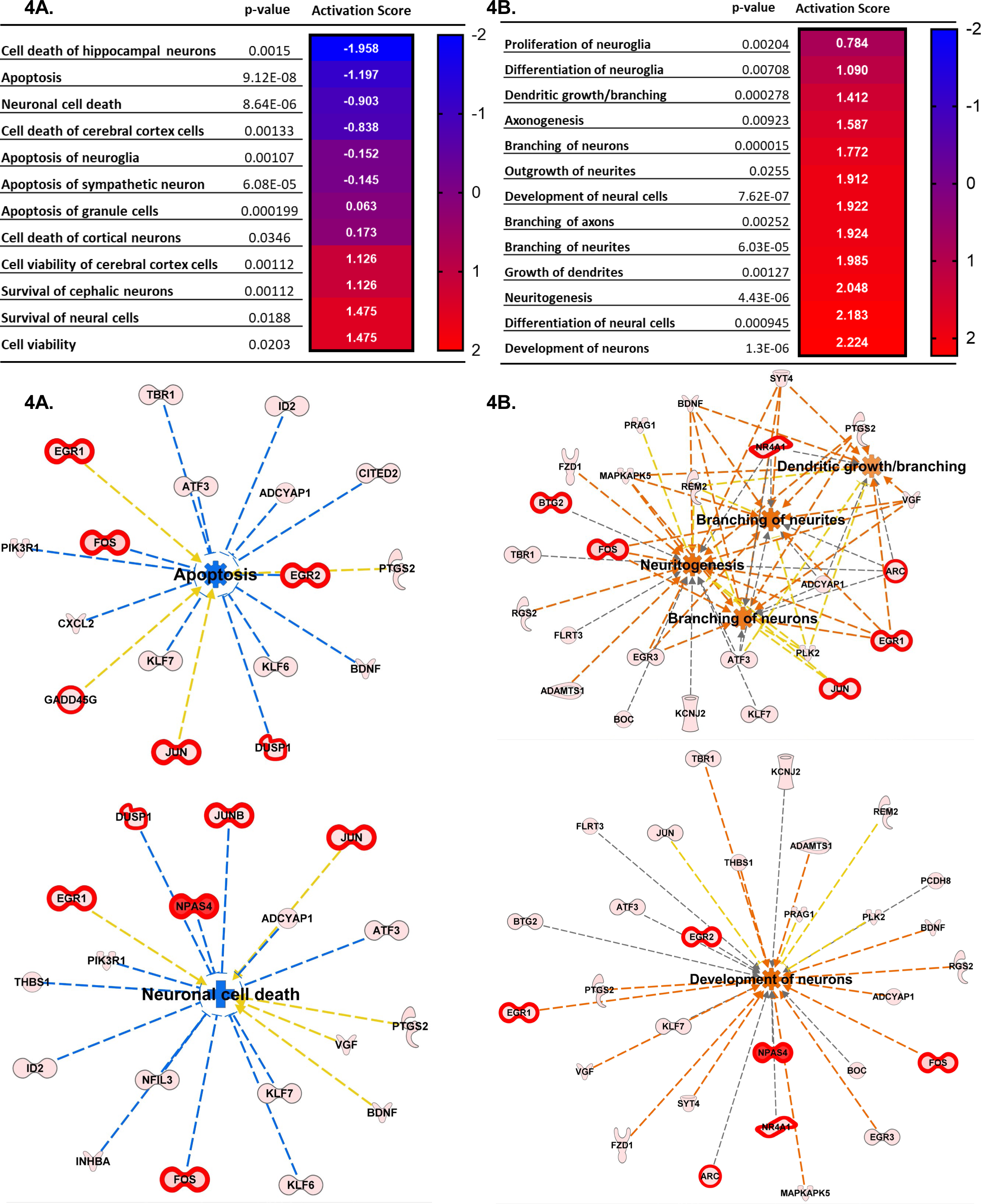
Top Cellular functions generated by IPA analysis of DEGs upregulated in SD animals vs Contralateral. A) Neural death and survival pathways. B) Cellular proliferation and growth. Top cellular functions generated by IPA analysis of DEGs upregulated in SD animals vs sham. The heatmaps show the p value and the activation score for each pathway. A’) and B’) Graphical representations of the networks and the genes involved in A and B. The intensity of the color red for each molecule is proportional to the FC of the gene. Orange arrows indicate a positive control of the gene on the related pathway, and the pathway’s name highlighted in orange indicates its activation. Genes with FDR< 0.01 are also highlighted with thicker red outlines. Apoptosis z score -2.003, neuronal cell death z score -1.974, density of spines z score +1.165, branching of neurons z score +1.913, dendritic growth/branching z score +1.090, and development of neurons z score +2.286.

**Table 2.** Predictions of SD-induced changes associated nervous system disease states and functions, in SD ipsilateral vs contralateral cortex.

| **Disease and Disorders** | **p-value range** | **# Molecules** |
| --- | --- | --- |
| Neurological Diseases | 4.56E-02 - 2.49E-08 | 73 |
| Organismal Injury and Abnormalities | 4.56E-02 - 1.91E-08 | 73 |
| Psychological Disorders | 2.85E-02 - 1.91E-08 | 35 |
| Hereditary Disorders | 1.54E-02 - 1.91E-08 | 21 |
| Skeletal and Muscular Disorders | 7.75E-03 - 6.30E-08 | 20 |
| **Physiological System Development and Functions** | **p-value range** | **# Molecules** |
| Nervous System Development and Function | 4.56E-02 - 7.04E-07 | 61 |
| Tissue Morphology | 4.56E-02 - 1.07E-09 | 39 |
| Tissue Development | 4.56E-02 - 1.03E-06 | 49 |
| Organismal Development | 4.56E-02 - 5.03E-06 | 44 |
| Organ Morphology | 4.56E-02 - 5.96E-05 | 16 |

### 3.4 SD target genes control cell differentiation and proliferation, synaptic plasticity, and stress responses

To validate the predicted activation of neuroprotective pathways, we analyzed the expression levels of a subset of SD target genes by qRT-PCR. The genes selected for validation where based on the IPA analysis; indeed, enrichment pathways analysis was used to identify genes functionally related to SD target genes that were not found significant in DE analysis. Based on the IPA results above, we focused on genes regulating 3 cellular functions: 1) cell differentiation and proliferation, 2) synaptic plasticity and 3) stress responses. qPCR was performed on the same RNA that was subjected to the RNAseq analyses, as well as 3 additional animals for each group. As shown in Figure 5, multiple SDs resulted in significant increases in the expression of genes activating cell proliferation and differentiation such as BNDF, DUPS6, FOS and JUNB (Figure 5). Considering the predicted activation of neurons and long-term potentiation, as well the increased axonogenesis, growth of neurites, and axon and neurites branching, we then analyzed the expression of genes known for their regulation of synaptic plasticity. As shown in Figure 5 we found significantly increased expression of Homer1a and ARC. Finally, we examined levels of PTGS2, NR4A1, and EGR2 as genes regulating stress and inflammatory responses, showing a significant upregulation.

**Figure 5.**
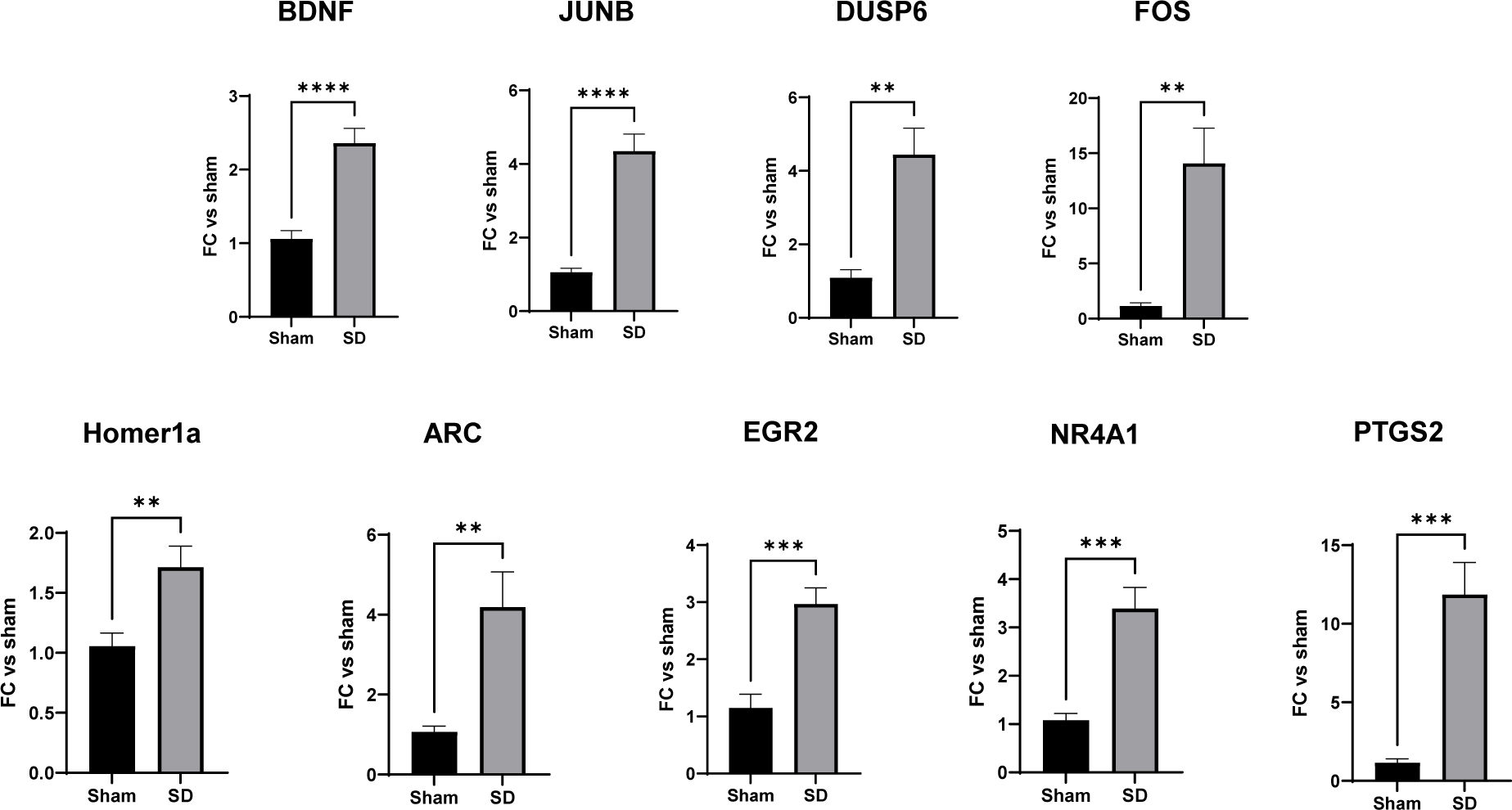
SD target genes control cell differentiation and proliferation, synaptic plasticity, and stress responses. RT-qPCR validation of SD target genes identified by RNAseq and enrichment analysis. mRNA levels of the target genes were measured as described in Material and Methods. Data were analyzed by unpaired t-test, followed by Welch’s correction and are represented as mean ± SEM; p<0.05*, **p<0.01, ***p<0.001; n=6.

### 3.5 Target genes showed differential expression depending on the distance from the SD initiation site

Lastly, we tested whether the levels of gene expression varied with distance from the SD initiation site. In this set of studies, KCl was used as the stimulus, as the site of initiation is clearly restricted to small zone of the application of KCl through the burr hole. Thus, two hours after onset of the initial SD, RNA was extracted from either total contralateral cortex, or from 3 different regions of the ipsilateral hemisphere: 1) SD initiation site; 2) intermediate site, and 3) remote site (see Methods and **Supplementary** Figure 3). By qRT-PCR we analyzed the levels of the previously identified target genes (see section 3 above) and compared these to the levels in the contralateral hemisphere of SHAM animals. For the majority of the genes examined (BDNF, DUSP6, PTGS2 ARC, and Homer 1a), highest expression was detected in the intermediate area (>3mm from the initiation site; Figure 6) while FOS and JUNB genes showed significantly increased expression in the remote site (>6 mm from the initiation site). Interestingly, when analyzing Homer1a levels, we found a decrease in their expression at the SD initiation site (Figure 6) when compared to the contralateral and intermediate sites (Figure 6). The difference between the contralateral and remote sites for NR4A1 did not reach statistical significance (p value=0.09) and EGR2 did not show differential expression across regions (**Supplementary** Figure 3).

**Figure 6.**
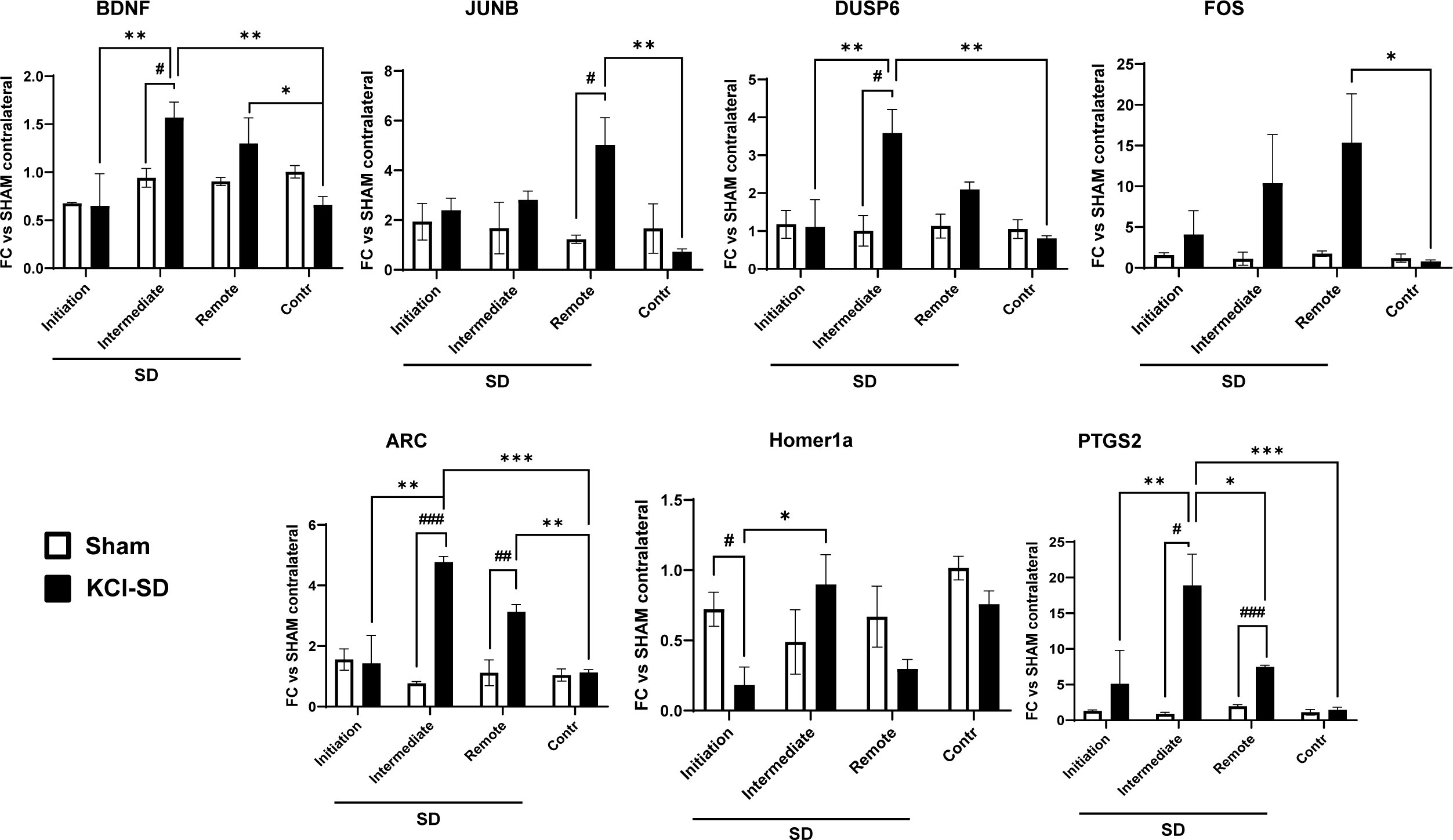
Target genes showed differential expression depending on the distance from the SD initiation site. mRNA levels of the target genes were measured as described in Material and Methods. Specifically, we determined the mRNA levels in 4 different regions: 1) SD initiation site, 2) SD intermediate site, 3) SD remote site, and 4) the contralateral hemisphere. Data were analyzed by 2-way ANOVA followed by Tukey’s Multiple Comparison test, and are represented as mean ± SEM; p<0.05*, **p<0.01, ***p<0.001; and unpaired t-test, followed by Welch’s correction and are represented as mean ± SEM; #p<0.05, ##p<0.01, ###p<0.001; n=3.

## 4 Discussion

This study is the first unbiased analysis of whole transcriptome changes in healthy brain tissue following repetitive spreading depolarization (SD). The main findings are that SD produces a spectrum of gene expression changes that are more diverse than previously thought and which, based on enrichment pathway analysis, likely include multiple molecular networks that promote neuronal survival, recovery and repair. The expression changes were most prominent in areas of SD propagation, implying that it is the depolarization wave itself that produces the changes rather than areas involved in SD initiation. We also show that the contralateral hemisphere, which is not directly involved in the SD, can serve as a good control tissue for DE analyses, even though changes in functional connectivity likely occur during repetitive SD waves. Our findings provide a basis for more consideration of a wider and novel range of consequences of SD, in addition to the well-studied acute functional disruption or damage to injured tissues.

SD is a profound activation of brain tissue that leads to depolarization in the order of minutes, and cellular loading of Na^+^, Ca^2+^ (Somjen, 2001; Hartings et al., 2017a), and activation of transcription factors (Ikeda et al., 1994; Sosthenes et al., 2019). In our study we confirmed that expression of immediate early genes is strongly upregulated, including that of established markers of neuronal activity such as FOS, JUNB and ARC (Ikeda et al., 1994; Hermann et al., 1999). SD is a global event that appears to involve all neurons and glia in its path (Somjen, 2001), and thus these markers of neuronal activation are expected to be uniformly activated in these cell types after SDs.

While RNAseq is a powerful tool for identifying genome-wide transcriptional alterations, standard differential gene expression analyses do not themselves leverage known biological gene network relationships. Using enrichment pathway analysis tools, we identified several additional key SD-activated molecules inferred as participants in larger functional networks. .

Increases in BDNF expression are among the most studied and strongly implicated in ischemic preconditioning and potentially plasticity after SD. BDNF increases were seen here as well. Consistent with previous work (Kokaia et al., 1993; Matsushima et al., 1998; Rangel et al., 2001; Yanamoto et al., 2004), BDNF was one of the genes significantly increased by both KCl and optogenetic stimuli (Figure 5 **and 6**) and was identified in SD-induced pathways related to neuronal differentiation: neuritogenesis, branching of axon sand neurites, and development of neurons (Figure 2 **and 4**). A surprising finding was the extent of predicted pathways involved in neuronal plasticity and regeneration. For example ARC mediates AMPA receptor endocytosis and is required for protein synthesis dependent LTP and memory formation (Chowdhury et al., 2006). Similarly, Homer1 scaffolding protein controls the insertion of postsynaptic glutamate receptors (Ango et al., 2002; Luo et al., 2012). Homer1a is increased by synaptic activity through the MAP kinase pathway. This type of regulation creates a negative feedback loop in which Homer1a inhibits Homer1b/c ability to scaffold and diminishes synaptic strength by modulation of dendritic spine morphology and AMPA/NMDA receptor activity (Sato et al., 2001; Bottai et al., 2002). This loop has been associated with TBI and epilepsy, particularly, and is thought to block postsynaptic Ca^2+^-mediated excitotoxicity (Cavarsan et al., 2012; Luo et al., 2014). Consistent with these observations, prior work in brain slices and rodent cortex has reported LTP-like synaptic strengthening following SD (Footitt and Newberry, 1998; Berger et al., 2008). In addition previous studies have shown that SD leads to synaptic strengthening (Footitt and Newberry, 1998; Berger et al., 2008; Theriot et al., 2012) as well as adult neurogenesis (Urbach et al., 2017) and preconditioning (Matsushima et al., 1998; Yanamoto et al., 2004; Shen et al., 2016). It will be of interest to determine whether the time course of associated gene expression changes correspond to variations in physiological recordings and how the expression of these genes change under different metabolic conditions surrounding ischemic infarcts.

Consistent with previous reports, either from gene array studies or targeted expression analysis, markers of inflammation were also upregulated (Urbach et al., 2006; Sintas et al., 2017). For example, our results are consistent with prior reports of robust increases of the pro-inflammatory mediator COX2 (Kaido et al., 2012; Takizawa et al., 2020; Volobueva et al., 2023). We did not see significant increases in some other inflammatory mediators that had been previously reported (e.g., TNF-α, IL-1β, IL6), perhaps because the time course for peak increases in these mediators is later than the 2-hour time point used in the present study (Takizawa et al., 2020). This time point was chosen based on 1) prior expression analysis confirming peak increases at 2 hours for most genes of interest (Kokaia et al., 1993; Karikó et al., 1998; Rangel et al., 2001) and 2) clinical recordings establishing clusters of SDs (usually defined as three or more events within 2 hours) as being clinically significant (Dreier et al., 2006; Hartings et al., 2020). Prior work (Takizawa et al., 2020) has shown that some inflammatory mediators can be detected following a single SD (measured at 1 hour), while others peak at 24 h (Matsushima et al., 1998; Jander et al., 2001; Volobueva et al., 2023). It remains to be determined which of the genes seen here may peak earlier or at later times points.

A second surprising finding came from pathway analysis of genes associated with neuronal injury or apoptotic cell death. The pathways analysis indicated a downregulation of these pathways, based on differential expression of molecular components of cell viability or apoptotic pathways such as: BDNF, FOS, EGR1 and 2, ATF3, NAPS4, CXCL2, and MAPKs (**Figure 2 and 4**). This raises the possibility that the extreme metabolic challenge of SD may be capable of promoting cell survival by decreasing cell death pathways. This does not mean that SDs are not causing neuronal injury – it means instead that SD is likely able to kill neurons due to extreme metabolic depletion, but in neurons that are able to recover from that acute insult, gene expression changes favor survival, including downregulation of cell death pathways. While further studies will be required to test if SDs is sufficient for a positive outcome after stroke or brain injury; this work shows that in tissues with sufficient perfusion, SD itself can induce the expression of neuroprotective pathways.

Another important experimental consideration that is somewhat unique for SD studies, is the choice of control tissue. SDs can propagate throughout a hemisphere but generally do not cross the midline and invade the contralateral hemisphere. For this reason, the contralateral hemisphere is often used as a control tissue for DEG studies (Shen and Gundlach, 1998; Wiggins et al., 2003; Volobueva et al., 2023), with the assumption that it is unaffected by SD. However, a well-established consequence of SD is a spreading depression of synaptic transmission throughout the affected cortex and, because of the connectivity between hemispheres (Leao, 1944; Vinogradova et al., 2021), an abrupt silencing of such large territory is expected to have consequences on circuit activity throughout the brain. We therefore tested whether the results of DEG analysis were different if the contralateral hemisphere was used instead of different animals in which no SD was generated (shams, as used in Figures 3 **and 4**). Interestingly, there was a very similar (although not identical) set of DEGs in both control tissues, implying differences in connectivity only cause minor differences in expression in the contralateral hemisphere compared with the changes produced by propagation of SD through the ipsilateral cortex. This suggests that the benefits of using control tissue from the same animal (to reduce inter-animal variability) may outweigh small changes in state for most SD studies.

Gene expression studies of SD have specific experimental challenges, related to the unusually extreme global disruptions produced by the events. Firstly, experimentally-induced SDs require coordinated depolarization of a volume of brain tissue that can be accomplished by a range of methods. Two commonly used methods (focal application of KCl and optogenetic stimulation) of SD were used here and combined, to increase generalizability of results. We then tested whether a microdissection and qPCR would show differences in propagation areas, as compared with the initiation sites. Interestingly, the largest expression changes were observed in the propagation regions (Figure 6), despite the fact that depolarization may be most intense at the initiation site. Factors released by SD or intracellular pathways activated by the combination of factors released at the wave front may be responsible. It also suggests that any injury or other damage produced at an initiation site was not the driver of changes observed in the study. Spatial genomics approaches (Ibarra-Lopez et al., 2022) will be useful to further investigate the distribution of changes of larger gene sets with distance from SD initiation, and with respect to differential vascular perfusion, for example.

In conclusion, these findings provide a basis for considering long-term changes due to SD that can influence outcomes in a range of neurological disorders. While SDs can clearly be damaging to vulnerable brain tissue, our work shows the changes in gene expression may also promote neuronal survival in some areas. If the predicted pathways involved in neuronal differentiation and growth are manifest in surviving peri-infarct tissues, for example, then there may be benefit to preserving these effects while preventing deleterious consequences of SD. These results also emphasize the importance of experimental design for SD studies. Finally, SDs that are experimentally generated outside a zone of vulnerable tissue may have the ability to activate pro-survival pathways predicted in the current study and thereby contribute to apparent paradoxical results in experimental models of stroke. Further studies will be essential to determine whether SD induction of neuronal survival pathways are sufficient to improve recovery and outcomes in stroke and brain injury.

## 5 Data Availability Statement

The data presented in this publication have been deposited in NCBI’s Gene Expression Omnibus (Edgar et al., 2002) and are accessible through GEO Series accession number GSE214623 (https://www.ncbi.nlm.nih.gov/geo/query/acc.cgi?acc=GSE214623).

## 7 Author Contributions

MD designed and performed the experiments, analysed the data, and wrote the manuscript together with JW, NPB, RM and DL. CWS supervised the project and was in charge of the overall planning and data analysis. All authors discussed the results and contributed to the final manuscript.

## 9 Acknowledgments

We thank the Animal Research Facility (ARF) of the Health Science Center (HSC), University of New Mexico School of Medicine. We also thank the UNM Preclinical Core of the Center from Brain Recovery and Repair (University of New Mexico) for technical support.

## 10 Ethics Statement

The animal study was reviewed and approved by the University of New Mexico Health Sciences Center Institutional Animal Care and Use Committee.

---

**Sup Figure 1.**
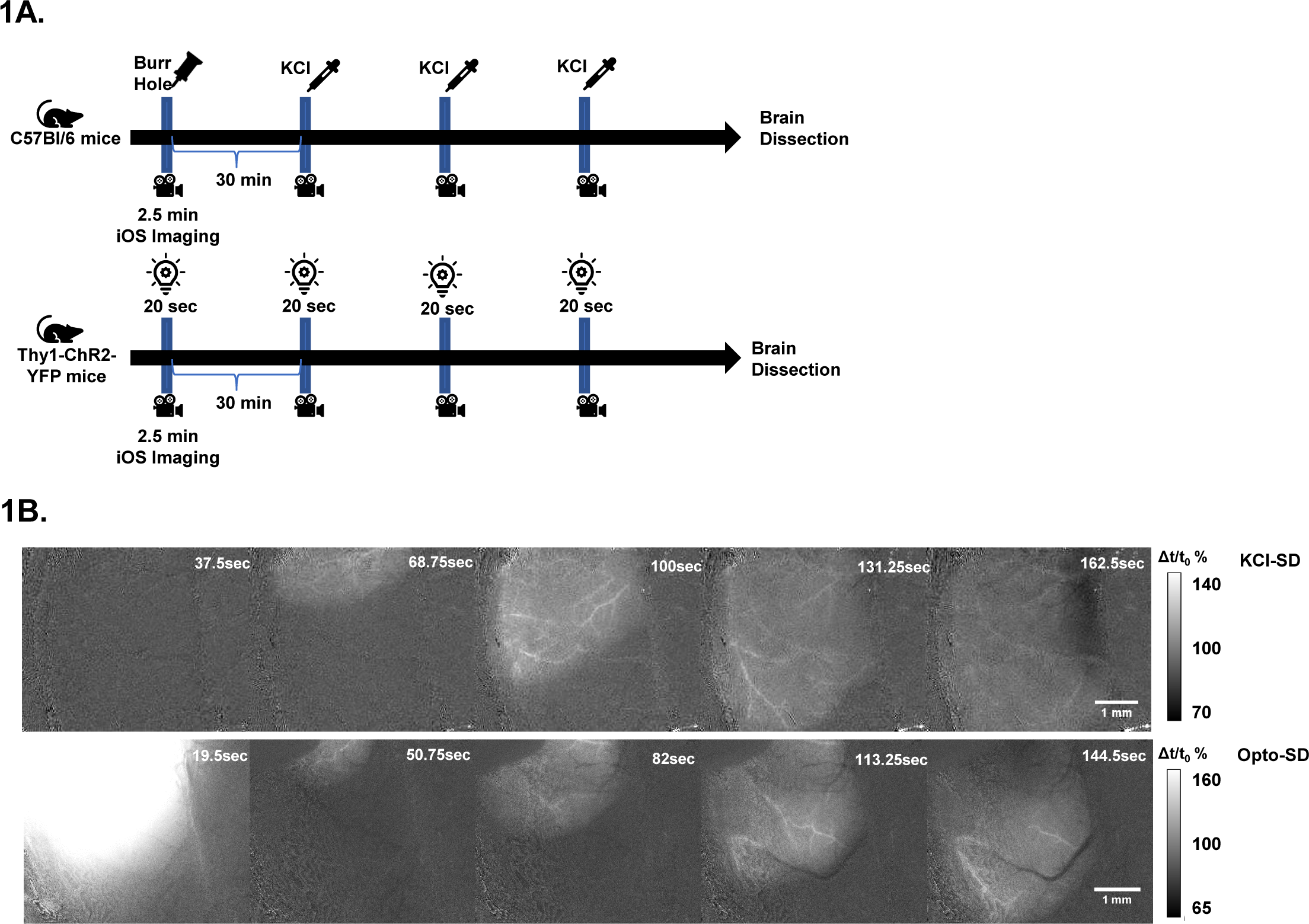

**Sup Figure 2.**
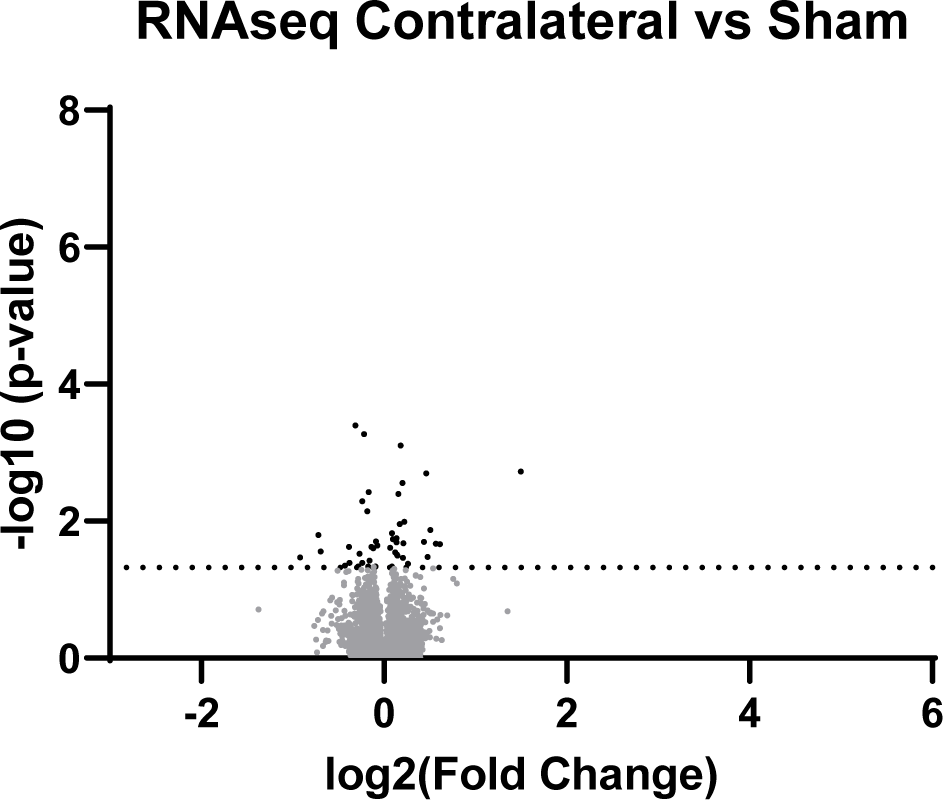

## 11 References

Aiba, I., Carlson, A. P., Sheline, C. T., and William, S. C. (2012). Synaptic release and extracellular actions of zn 2+ limit propagation of spreading depression and related events in vitro and in vivo. J. Neurophysiol. 107, 1032–1041. doi: 10.1152/jn.00453.2011.

Ango, F., Robbe, D., Tu, J. C., Xiao, B., Worley, P. F., Pin, J.-P., et al. (2002). Homer-dependent cell surface expression of metabotropic glutamate receptor type 5 in neurons. Mol. Cell. Neurosci. 20, 323–9. Available at: http://www.ncbi.nlm.nih.gov/pubmed/12093163 [Accessed August 13, 2019].

Arenkiel, B. R., Peca, J., Davison, I. G., Feliciano, C., Deisseroth, K., Augustine, G. J. J., et al. (2007). In vivo light-induced activation of neural circuitry in transgenic mice expressing channelrhodopsin-2. Neuron 54, 205–218. doi: 10.1016/J.NEURON.2007.03.005.

Ayata, C., and Lauritzen, M. (2015). Spreading Depression, Spreading Depolarizations, and the Cerebral Vasculature. Physiol. Rev. 95, 953–993. doi: 10.1152/PHYSREV.00027.2014.

Berger, M., Speckmann, E. J., Pape, H. C., and Gorji, A. (2008). Spreading depression enhances human neocortical excitability in vitro. Cephalalgia 28, 558–562. doi: 10.1111/J.1468-2982.2008.01556.X.

Bottai, D., Guzowski, J. F., Schwarz, M. K., Kang, S. H., Xiao, B., Lanahan, A., et al. (2002). Synaptic Activity-Induced Conversion of Intronic to Exonic Sequence in Homer 1 Immediate Early Gene Expression. J. Neurosci. 22, 167. doi: 10.1523/JNEUROSCI.22-01-00167.2002.

Carlson, A. P., Abbas, M., Alunday, R. L., Qeadan, F., and William Shuttleworth, C. (2018). Spreading depolarization in acute brain injury inhibited by ketamine: a prospective, randomized, multiple crossover trial. J. Neurosurg. 130, 1513–1519. doi: 10.3171/2017.12.JNS171665.

Cavarsan, C. F., Tescarollo, F., Tesone-Coelho, C., Morais, R. L. T., Motta, F. L. T., Blanco, M. M., et al. (2012). Pilocarpine-induced status epilepticus increases Homer1a and changes mGluR5 expression. Epilepsy Res. 101, 253–260. doi: 10.1016/J.EPLEPSYRES.2012.04.011.

Charles, A., and Brennan, K. C. (2009). Cortical spreading depression-new insights and persistent questions. Cephalalgia 29, 1115–1124. doi: 10.1111/J.1468-2982.2009.01983.X.

Chowdhury, S., Shepherd, J. D., Okuno, H., Lyford, G., Petralia, R. S., Plath, N., et al. (2006). Arc Interacts with the Endocytic Machinery to Regulate AMPA Receptor Trafficking. Neuron 52, 445. doi: 10.1016/J.NEURON.2006.08.033.

Dietrich, W. D., Truettner, J., Prado, R., Stagliano, N. E., Zhao, W., Busto, R., et al. (2000). Thromboembolic events lead to cortical spreading depression and expression of c-fos, brain-derived neurotrophic factor, glial fibrillary acidic protein, and heat shock protein 70 mRNA in rats. J. Cereb. Blood Flow Metab. 20, 103–11. doi: 10.1097/00004647-200001000-00014.

Dreier, J. P. (2011). The role of spreading depression, spreading depolarization and spreading ischemia in neurological disease. Nat. Med. 17, 439–447. doi: 10.1038/nm.2333.

Dreier, J. P., Fabricius, M., Ayata, C., Sakowitz, O. W., William Shuttleworth, C., Dohmen, C., et al. (2017). Recording, analysis, and interpretation of spreading depolarizations in neurointensive care: Review and recommendations of the COSBID research group. J. Cereb. Blood Flow Metab. 37, 1595–1625. doi: 10.1177/0271678X16654496.

Dreier, J. P., Woitzik, J., Fabricius, M., Bhatia, R., Major, S., Drenckhahn, C., et al. (2006). Delayed ischaemic neurological deficits after subarachnoid haemorrhage are associated with clusters of spreading depolarizations. Brain 129, 3224–3237. doi: 10.1093/BRAIN/AWL297.

Edgar, R., Domrachev, M., and Lash, A. E. (2002). Gene Expression Omnibus: NCBI gene expression and hybridization array data repository. Nucleic Acids Res. 30, 207–210. doi: 10.1093/NAR/30.1.207.

Eising, E., Shyti, R., ’t Hoen, P. A. C., Vijfhuizen, L. S., Huisman, S. M. H., Broos, L. A. M., et al. (2017). Cortical Spreading Depression Causes Unique Dysregulation of Inflammatory Pathways in a Transgenic Mouse Model of Migraine. Mol. Neurobiol. 54, 2986–2996. doi: 10.1007/S12035-015-9681-5.

Footitt, D. R., and Newberry, N. R. (1998). Cortical spreading depression induces an LTP-like effect in rat neocortex in vitro. Brain Res. 781, 339–342. doi: 10.1016/S0006-8993(97)01359-0.

Ge, S. X., Son, E. W., and Yao, R. (2018). iDEP: An integrated web application for differential expression and pathway analysis of RNA-Seq data. BMC Bioinformatics 19, 1–24. doi: 10.1186/s12859-018-2486-6.

Hartings, J. A., Andaluz, N., Bullock, M. R., Hinzman, J. M., Mathern, B., Pahl, C., et al. (2020). Prognostic Value of Spreading Depolarizations in Patients With Severe Traumatic Brain Injury. JAMA Neurol. 77, 489–499. doi: 10.1001/JAMANEUROL.2019.4476.

Hartings, J. A., Li, C., Hinzman, J. M., Shuttleworth, C. W., Ernst, G. L., Dreier, J. P., et al. (2017a). Direct current electrocorticography for clinical neuromonitoring of spreading depolarizations. J. Cereb. Blood Flow Metab. 37, 1857–1870. doi: 10.1177/0271678X16653135.

Hartings, J. A., Shuttleworth, C. W., Kirov, S. A., Ayata, C., Hinzman, J. M., Foreman, B., et al. (2017b). The continuum of spreading depolarizations in acute cortical lesion development: Examining Leão’s legacy. J. Cereb. Blood Flow Metab. 37, 1571–1594. doi: 10.1177/0271678X16654495.

Helbok, R., Hartings, J. A., Schiefecker, A., Balança, B., Jewel, S., Foreman, B., et al. (2020). What Should a Clinician Do When Spreading Depolarizations are Observed in a Patient? Neurocrit. Care 32, 306–310. doi: 10.1007/s12028-019-00777-6.

Hermann, D. M., Mies, G., and Hossmann, K. A. (1999). Expression of c-fos, junB, c-jun, MKP-1 and hsp72 following traumatic neocortical lesions in rats--relation to spreading depression. Neuroscience 88, 599–608. doi: 10.1016/s0306-4522(98)00249-8.

Hinzman, J. M., DiNapoli, V. A., Mahoney, E. J., Gerhardt, G. A., and Hartings, J. A. (2015). Spreading depolarizations mediate excitotoxicity in the development of acute cortical lesions. Exp. Neurol. 267, 243–253. doi: 10.1016/j.expneurol.2015.03.014.

Ibarra-Lopez, V., Jayakar, S., Yang, Y. A., Martin, C., Modrusan, Z., and Rost, S. (2022). Spatial Profiling of Protein and RNA Expression in Tissue: An Approach to Fine-tune Virtual Microdissection. J. Vis. Exp. doi: 10.3791/62651.

Ikeda, J., Nakajima, T., Osborne, O. C., Mies, G., and Nowak, T. S. (1994). Coexpression of c-fos and hsp70 mRNAs in gerbil brain after ischemia: Induction threshold, distribution and time course evaluated by in situ hybridization. Mol. Brain Res. 26, 249–258. doi: 10.1016/0169-328X(94)90097-3.

Jander, S., Schroeter, M., Peters, O., Witte, O. W., and Stoll, G. (2001). Cortical spreading depression induces proinflammatory cytokine gene expression in the rat brain. J. Cereb. Blood Flow Metab. 21, 218–225. doi: 10.1097/00004647-200103000-00005.

Kaido, Kempski, Heimann, Heers, and Bartsch (2012). Cluster analysis of mRNA expression levels identifies multiple sequential patterns following focal cerebral ischemia. Turk. Neurosurg. 22. doi: 10.5137/1019-5149.JTN.5523-11.0.

Karikó, K., Harris, V. A., Rangel, Y., Duvall, M. E., and Welsh, F. A. (1998). Effect of cortical spreading depression on the levels of mRNA coding for putative neuroprotective proteins in rat brain. J. Cereb. Blood Flow Metab. 18, 1308–1315. doi: 10.1097/00004647-199812000-00005.

Kokaia, Z., Gidö, G., Ringstedt, T., Bengzon, J., Kokaia, M., Siesjö, B. K., et al. (1993). Rapid increase of BDNF mRNA levels in cortical neurons following spreading depression: regulation by glutamatergic mechanisms independent of seizure activity. Mol. Brain Res. 19, 277–286. doi: 10.1016/0169-328X(93)90126-A.

Krämer, A., Green, J., Pollard, J., and Tugendreich, S. (2014). Causal analysis approaches in Ingenuity Pathway Analysis. Bioinformatics 30, 523–530. doi: 10.1093/BIOINFORMATICS/BTT703.

Lauritzen, M., Dreier, J. P., Fabricius, M., Hartings, J. A., Graf, R., and Strong, A. J. (2011). Clinical relevance of cortical spreading depression in neurological disorders: Migraine, malignant stroke, subarachnoid and intracranial hemorrhage, and traumatic brain injury. J. Cereb. Blood Flow Metab. 31, 17–35. doi: 10.1038/jcbfm.2010.191.

Leao, A. A. P. (1944). SPREADING DEPRESSION OF ACTIVITY IN THE CEREBRAL CORTEX. J. Neurophysiol. 7, 359–390. doi: 10.1152/jn.1944.7.6.359.

Lindquist, B. E., and Shuttleworth, C. W. (2014). Spreading depolarization-induced adenosine accumulation reflects metabolic status in vitro and in vivo. J. Cereb. Blood Flow Metab. 34, 1779–1790. doi: 10.1038/jcbfm.2014.146.

Livak, K. J., and Schmittgen, T. D. (2001). Analysis of relative gene expression data using real-time quantitative PCR and the 2-ΔΔCT method. Methods 25, 402–408. doi: 10.1006/meth.2001.1262.

Luo, P., Chen, T., Zhao, Y., Zhang, L., Yang, Y., Liu, W., et al. (2014). Postsynaptic scaffold protein Homer 1a protects against traumatic brain injury via regulating group I metabotropic glutamate receptors. Cell Death Dis. 5. doi: 10.1038/CDDIS.2014.116.

Luo, P., Li, X., Fei, Z., and Poon, W. (2012). Scaffold protein Homer 1: Implications for neurological diseases. Neurochem. Int. 61, 731–738. doi: 10.1016/j.neuint.2012.06.014.

Matsushima, K., Hogan, M. J., and Hakim, A. M. (1996). Cortical spreading depression protects against subsequent focal cerebral ischemia in rats. J. Cereb. Blood Flow Metab. 16, 221–226. doi: 10.1097/00004647-199603000-00006.

Matsushima, K., Schmidt-Kastner, R., Hogan, M. J., and Hakim, A. M. (1998). Cortical spreading depression activates trophic factor expression in neurons and astrocytes and protects against subsequent focal brain ischemia. Brain Res. 807, 47–60. doi: 10.1016/s0006-8993(98)00716-1.

Nedergaard, M., and Hansen, A. J. (1988). Spreading depression is not associated with neuronal injury in the normal brain. Brain Res. 449, 395–398. doi: 10.1016/0006-8993(88)91062-1.

Pertea, M., Kim, D., Pertea, G. M., Leek, J. T., and Salzberg, S. L. (2016). Transcript-level expression analysis of RNA-seq experiments with HISAT, StringTie and Ballgown. Nat. Protoc. 2016 *119* 11, 1650–1667. doi: 10.1038/nprot.2016.095.

Pietrobon, D., and Moskowitz, M. A. (2014). Chaos and commotion in the wake of cortical spreading depression and spreading depolarizations. Nat. Rev. Neurosci. 15, 379–393. doi: 10.1038/NRN3770.

Rangel, Y. M., Karikó, K., Harris, V. A., Duvall, M. E., and Welsh, F. A. (2001). Dose-dependent induction of mRNAs encoding brain-derived neurotrophic factor and heat-shock protein-72 after cortical spreading depression in the rat. Mol. Brain Res. 88, 103–112. doi: 10.1016/S0169-328X(01)00037-7.

Sato, M., Suzuki, K., and Nakanishi, S. (2001). NMDA receptor stimulation and brain-derived neurotrophic factor upregulate homer 1a mRNA via the mitogen-activated protein kinase cascade in cultured cerebellar granule cells. J. Neurosci. 21, 3797–3805. doi: 10.1523/JNEUROSCI.21-11-03797.2001.

Shen, P. J., and Gundlach, A. L. (1998). Differential spatiotemporal alterations in adrenoceptor mRNAs and binding sites in cerebral cortex following spreading depression: Selective and prolonged up-regulation of α(1B)-adrenoceptors. Exp. Neurol. 154, 612–627. doi: 10.1006/exnr.1998.6915.

Shen, P. P., Hou, S., Ma, D., Zhao, M. M., Zhu, M. Q., Zhang, J. D., et al. (2016). Cortical spreading depression-induced preconditioning in the brain. Neural Regen. Res. 11, 1857–1864. doi: 10.4103/1673-5374.194759.

Sintas, C., Fernàndez-Castillo, N., Vila-Pueyo, M., Pozo-Rosich, P., Macaya, A., and Cormand, B. (2017). Transcriptomic Changes in Rat Cortex and Brainstem After Cortical Spreading Depression With or Without Pretreatment With Migraine Prophylactic Drugs. J. Pain 18, 366–375. doi: 10.1016/j.jpain.2016.11.007.

Somjen, G. G. (2001). Mechanisms of spreading depression and hypoxic spreading depression-like depolarization. Physiol. Rev. 81, 1065–1096. doi: 10.1152/PHYSREV.2001.81.3.1065.

Sosthenes, M. C. K., Diniz, D. G., Roodselaar, J., Abadie-Guedes, R., de Siqueira Mendes, F. de C. C., Fernandes, T. N., et al. (2019). Stereological Analysis of Early Gene Expression Using Egr-1 Immunolabeling After Spreading Depression in the Rat Somatosensory Cortex. Front.Neurosci. 13, 1020. doi: 10.3389/fnins.2019.01020.

Takizawa, T., Qin, T., Morais, A. L. de, Sugimoto, K., Chung, J. Y., Morsett, L., et al. (2020). Non-invasively triggered spreading depolarizations induce a rapidpro-inflammatory response in cerebral cortex. J. Cereb. Blood Flow Metab. 40, 1117. doi: 10.1177/0271678X19859381.

Theriot, J. J., Toga, A. W., Prakash, N., Sungtaek Ju, Y., and Brennan, K. C. (2012). Cortical sensory plasticity in a model of migraine with aura. J. Neurosci. 32, 15252–15261. doi: 10.1523/JNEUROSCI.2092-12.2012.

Urbach, A., Baum, E., Braun, F., and Witte, O. W. (2017). Cortical spreading depolarization increases adult neurogenesis, and alters behavior and hippocampus-dependent memory in mice. J. Cereb. Blood Flow Metab. 37, 1776–1790. doi: 10.1177/0271678X16643736.

Urbach, A., Bruehl, C., and Witte, O. W. (2006). Microarray-based long-term detection of genes differentially expressed after cortical spreading depression. Eur. J. Neurosci. 24, 841–856. doi: 10.1111/j.1460-9568.2006.04862.x.

Vinogradova, L. V., Suleymanova, E. M., and Medvedeva, T. M. (2021). Transient loss of interhemispheric functional connectivity following unilateral cortical spreading depression in awake rats. Cephalalgia 41, 353–365. doi: 10.1177/0333102420970172.

Volobueva, M. N., Suleymanova, E. M., Smirnova, M. P., Bolshakov, A. P., and Vinogradova, L. V. (2023). A Single Episode of Cortical Spreading Depolarization Increases mRNA Levels of Proinflammatory Cytokines, Calcitonin Gene-Related Peptide and Pannexin-1 Channels in the Cerebral Cortex. Int. J. Mol. Sci. 24. doi: 10.3390/ijms24010085.

Wiggins, A. K., Shen, P.-J. J., and Gundlach, A. L. (2003). Atrial natriuretic peptide expression is increased in rat cerebral cortex following spreading depression: possible contribution to sd-induced neuroprotection. Neuroscience 118, 715–726. doi: 10.1016/s0306-4522(03)00006-x.

Yanamoto, H., Xue, J. H., Miyamoto, S., Nagata, I., Nakano, Y., Murao, K., et al. (2004). Spreading depression induces long-lasting brain protection against infarcted lesion development via BDNF gene-dependent mechanism. Brain Res. 1019, 178–188. doi: 10.1016/j.brainres.2004.05.105.

